# Dietary polyunsaturated fatty acid supply improves *Daphnia* performance during vertical migration

**DOI:** 10.1101/586156

**Authors:** Jana Isanta Navarro, Matthias Fromherz, Michelle Dietz, Bettina Zeis, Anke Schwarzenberger, Dominik Martin-Creuzburg

## Abstract

1. Diel vertical migration (DVM) is a common predator avoidance strategy of zooplankton. Migration to deeper water layers during the day to escape visually hunting predators most likely requires physiological adaptations to periodically changing temperatures. Polyunsaturated fatty acids (PUFA) are essential nutrients that play crucial roles in membrane temperature acclimation. Exposure to cold temperatures typically results in an increase in the relative abundance of PUFA in cell membranes and PUFA requirements of *Daphnia* have been shown to increase with decreasing temperatures.
2. To assess the significance of dietary PUFA for coping with temperature fluctuations experienced during DVM, we reared *Daphnia magna* at either constantly warm or fluctuating temperatures simulating DVM both with and without dietary PUFA supplementation.
3. We show that the well-known positive effect of dietary eicosapentaenoic acid (EPA) supplementation on offspring production and population growth rates of *Daphnia* is more pronounced at alternating temperatures than at constantly warm temperatures. Exposure to alternating temperatures caused modification in body PUFA concentrations and, consequently, increased lipid peroxidation. However, detrimental effects of lipid peroxidation were not evident.
4. Our data demonstrate that the capacity to cope with the distinct temperature fluctuations experienced during DVM increases with dietary EPA supplementation, suggesting that an adequate dietary PUFA supply is crucial especially for migrating *Daphnia* populations. A dietary deficiency in long-chain PUFA may thus severely constrain predator evasion, potentially resulting in increased mortality and cascading effects on lower trophic levels.

## Introduction

Diel vertical migration (DVM) has been recognized as a very efficient predator avoidance strategy of aquatic organisms, which is induced by chemical cues released from visually hunting predators and triggered by light availability (Siebeck 1960; Hays, 2003; Lampert, 2011). Species of the genus *Daphnia* are important model organisms for studying DVM (Rinke & Petzoldt, 2008; Lampert, 2011). Many *Daphnia* population are migrating to deeper water layers during the day to escape fish predation and thus are confronted with diurnal changes in temperature and food supply (Loose & Dawidowicz, 1994; Lampert, McCauley, & Manly, 2003). The amplitude of DVM can vary from a few up to 40 and more meters, depending on a variety of different factors, such as lake depth, light conditions and predation risk (Stich & Lampert, 1981; Dodson, 1990; von Elert & Pohnert 2000; Lampert, 2011). During DVM, *Daphnia* populations can experience temperature changes exceeding 15 °C (Stich & Lampert, 1981). Metabolic disadvantages associated with the exposure to cold temperatures have been identified as the primary cost of DVM, resulting in reduced growth and reproduction (Stich & Lampert, 1984; Lampert, 2011).

Exposure to cold temperature requires physiological adaptations, which have been poorly studied in regard to DVM. Restructuring the lipid composition of cell membranes is one important mechanism by which ectotherms adapt to changing temperatures. The homeoviscous adaptation hypothesis predicts that the proportion of unsaturated fatty acids in cell membranes increases with decreasing temperature to maintain vital membrane properties (Hazel & Williams, 1990; Hazel, 1995). Polyunsaturated fatty acids (PUFA) are essential nutrients for most animals; they are indispensable as structural components of cell membranes and as precursors for eicosanoids, signaling molecules acting primarily on reproduction and immunity (Stanley, 2000; Leonard, Pereira, Sprecher *et al*., 2004; Valentine & Valentine, 2004). The dietary supply with long-chain (>C18) PUFA, such as eicosapentaenoic acid (EPA), strongly affects growth and reproduction of *Daphnia* (von Elert, 2002; Becker & Boersma, 2003; Martin-Creuzburg, Wacker, & Basen, 2010). Dietary PUFA requirements of *Daphnia* have been shown to increase with decreasing temperature, suggesting that the dietary PUFA supply is crucial for coping with low temperatures (Sperfeld & Wacker, 2011; Martin-Creuzburg, Wacker, Ziese *et al*., 2012). Behavioral experiments suggest that *Daphnia* with access to dietary EPA are migrating to deeper water layers, presumably because of a higher PUFA-mediated capacity to cope with colder temperatures (Brzezinski & von Elert, 2015). The influence of fluctuating temperatures on PUFA requirements has not been studied yet. Despite the significance of PUFA for physiological processes, PUFA accumulation may also come along with costs because PUFA are particularly sensitive to lipid peroxidation, resulting in radical formation and oxidative stress (Yin, Xu, & Porter, 2011). Effects of temperature on lipid peroxidation and oxidative stress are controversial but most studies on ectotherms have found increasing oxidative stress with increasing temperature (Lushchak, 2011).

To assess the significance of dietary PUFA for coping with temperature fluctuations experienced during DVM, we reared *Daphnia magna* at either constantly warm or fluctuating temperatures simulating DVM both with and without dietary EPA supply, measured various life history traits, and analyzed the fatty acid composition of the animals. The thiobarbituric acid (TBA) assay was used to assess the effects of temperature and dietary PUFA supply on lipid peroxidation. We used *D. magna* as a model because this species is known to perform pronounced DVM if conditions are appropriate (Lampert, 2011) and is extensively used to study essential lipid requirements.

## Methods

### Cultivation of organisms

The experiment was performed with a *Daphnia magna* clone originally isolated from Grosser Binnensee, Germany (Lampert, 1991), which has been used previously to study DVM under laboratory conditions (von Elert & Pohnert, 2000; Brzezinski & von Elert, 2015). *Daphnia* were pre-raised at 20 °C in 1-L jars (10 individuals per jar) containing filtered lake water (<0.2 µm) and 2 mg C L^−1^ of the green alga *Acutodesmus obliquus* (formerly referred to as *Scenedesmus obliquus*; SAG 276-3a). *A. obliquus* was used as food also in the growth experiment because it lacks long-chain PUFA (i.e. >C18) and thus is of moderate food quality for *Daphnia* (von Elert, 2002, Martin-Creuzburg *et al*., 2012). *A. obliquus* was grown at 20 °C in 5-L batch cultures in Cyano medium (Jüttner, Leonhardt, & Möhren, 1983) and harvested in the late-exponential growth phase (illumination at 100 µmol quanta m^−2^ s^−1^). Food suspensions were prepared by centrifugation and resuspension in fresh medium. Carbon concentrations of the food suspensions were estimated from photometric light extinctions using previously established carbon-extinction regressions.

### Growth experiment

The experiment was conducted with third-clutch offspring born within 12 h. Neonates were placed individually in jars containing 80 ml of filtered (<0.2 µm) lake water and exposed to two different temperature regimes and two different food qualities in a full-factorial design. A subsample of 30 animals was taken from the isolated cohort of neonates for initial dry mass determination after freeze-drying. The jars were placed in a flow-through system in which they were constantly surrounded by temperature-controlled water of either constantly 20 °C or alternating 20 °C and 6 °C. The alternating temperature regime was set up to simulate the temperature changes that *Daphnia* potentially experience during DVM in deep, temperate lakes, i.e. eight hours at 20 °C, two hours for cooling down to 6 °C, 12 hours at 6 °C, and two hours for warming up to 20 °C again. All jars were provided with 2 mg C L^−1^ of *A. obliquus*. Food quality was manipulated by adding EPA-containing liposomes to the jars; liposomes prepared without adding EPA were used as a control supplement. Liposomes were prepared as described previously (Martin-Creuzburg, Sperfeld, & Wacker, 2009). Initially, each treatment consisted of 40 jars each containing one animal. Daphnids were transferred daily into new jars with freshly prepared food suspensions. At day six, 12 daphnids (jars) were subsampled, rinsed with ultra-pure water, transferred into pre-weighed aluminum boats, and stored at −80°C until they were freeze-dried and weighed (±0.1 µg) for dry mass determination and subsequent fatty acid and lipid peroxidation analysis. Fatty acids were analyzed as described in Martin-Creuzburg *et al*. (2010). Mass-specific juvenile somatic growth rates (g) were determined as the increase in dry mass from the beginning (M_0_) until day six of the experiment (M_t_) with time (t) expressed as age in days:

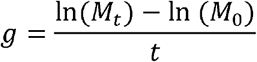

The remaining daphnids (28 replicates) were kept in their respective treatments until they released their third-clutch offspring. Population growth rates (r) were estimated iteratively using the Euler-Lotka equation:

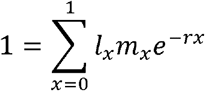

where l_x_ is the age-specific survivorship, m_x_ is the number of offspring at each reproduction cycle, and x is the age at reproduction in days (offspring production was recorded once a day, when the animals were transferred to new jars).

The concentration of malonic dialdehyde (MDA), an indicator of lipid peroxidation, was estimated using the thiobarbituric acid (TBA) assay (Ohkawa, Ohishi, & Yagi, 1979) and the data were expressed as TBA-reactive substances (TBARS). The concentration of TBARS was used as an approximation for the amount of damage caused by lipid peroxidation.

### Data analyses

Somatic growth rates, population growth rates, C18 body PUFA concentrations, and TMOP-equivalents were analysed using two-factorial ANOVAs followed by Tukey’s HSD post hoc tests (STATISTICA, StatSoft, Inc., 2011, Version 10.0.). Population growth rate data were log-transformed to meet ANOVA assumptions, i.e. homogeneity of variances. Age at first reproduction and cumulative counts of offspring were analysed using a generalized linear model (GLM) with log as the link function for quasi-Poisson errors to compensate for overdispersion (R Development Core Team, 2017). To compare the increase in somatic growth rates (g; d^−1^), offspring production (off; No.), and population growth rates (r; d^−1^) upon EPA supplementation between the two temperature regimes (Fig. 1, inserted bar graphs), the data were standardized as improvement (I) upon EPA supplementation relative to the values obtained on the unsupplemented food using the equation:

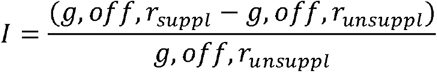

**Fig. 1:**
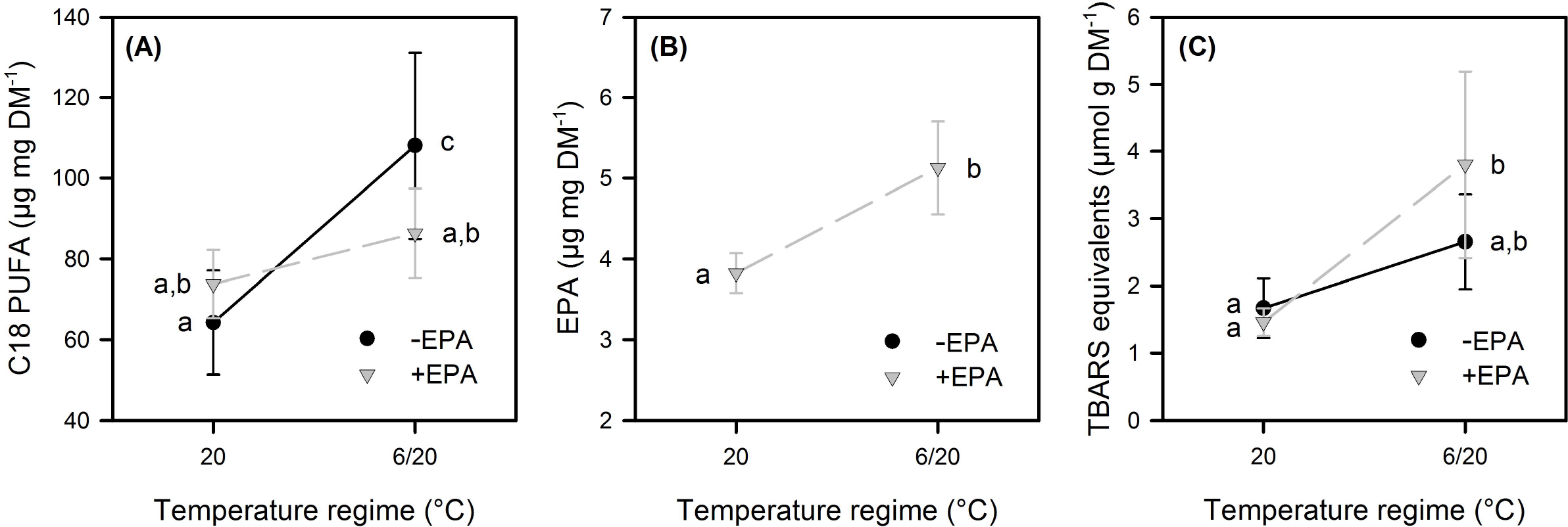
Juvenile somatic growth rates, age at first reproduction, cumulative numbers of offspring produced within the first three reproduction cycles, and population growth rates of *Daphnia magna* exposed to two different temperature regimes, i.e. either constantly 20°C or alternating 6 and 20°C. Data represent means ± standard deviation (n = 12 for somatic growth rates; n = 26-28 for offspring and population growth rates). Different letters indicate significant differences among treatments (Tukey’s HSD, P < 0.05). Inserted bar graphs depict the relative improvement in somatic growth rates, age at first reproduction, offspring production, and population growth rates upon EPA supplementation in the two temperature regimes.

## Results

Juvenile somatic growth rates were significantly affected by food and temperature (Fig. 1A, Table 1), i.e. they increased upon EPA supplementation in both temperature regimes and were significantly lower in the alternating temperature regime than in the constantly warm temperature regime. The increase in somatic growth rates upon EPA supplementation (Fig. 1A, inserted bar graph) and the significant interaction between food and temperature reflect that the positive effect of EPA supplementation was higher in the constantly warm temperature regime than in the alternating temperature regime.

**Table 1:** Results of the two-factorial ANOVA used to assess differences in juvenile somatic growth rates, population growth rates, C18 body PUFA concentrations, and TBARS-equivalents between food and temperature treatments (Fig. 1 and Fig. 2). For the age at first reproduction and the cumulative numbers of offspring produced in the first three reproduction cycles results of the generalized linear model are shown. Population growth rates were log-transformed to meet ANOVA assumptions. Asterisks (*) indicate significant factors and interactions.

|  | factor | df | *F* | *P(>F)* |
| --- | --- | --- | --- | --- |
| somatic growth rates | food | 1/44 | 70.7 | <0.001* |
|  | temperature | 1/44 | 1289.8 | <0.001* |
|  | food×temp | 1/44 | 11.9 | <0.001* |
| population growth rates | food | 1/104 | 196.0 | <0.001* |
|  | temperature | 1/104 | 4321.1 | <0.001* |
|  | food×temp | 1/104 | 74.6 | <0.001* |
| C18 PUFA | food | 1/9 | 0.6 | 0.48 |
|  | temperature | 1/9 | 11.7 | 0.008* |
|  | food×temp | 1/9 | 3.6 | 0.09 |
| TBARS equivalents | food | 1/11 | 1.2 | 0.30 |
|  | temperature | 1/11 | 15.1 | 0.003* |
|  | food×temp | 1/11 | 2.5 | 0.14 |
|  | factor | df | *Deviance* | *P(>X)* |
| cumulative offspring | food | 1/106 | 399.3 | <0.001* |
|  | temperature | 1/105 | 10.36 | 0.03 * |
|  | food×temp | 1/104 | 70.04 | <0.001* |
| age at first reproduction | food | 1/106 | 2.33 | <0.001* |
|  | temperature | 1/105 | 209.7 | <0.001* |
|  | food×temp | 1/104 | 1.54 | <0.001* |

The age at first reproduction was significantly affected by food and temperature (Fig. 1B, Table 1). Although EPA supplementation did not affect the age at first reproduction in the constantly warm temperature regime (on average 7.3 days in both food treatments) it resulted in a significantly younger age at first reproduction in the alternating temperature regime (on average 18.1 days without EPA, 15.7 days with EPA; Fig. 1B, Table 1).

The cumulative numbers of offspring were also significantly affected by food and temperature (Fig. 1C, Table 1). Without EPA supplementation, offspring numbers were significantly lower in the alternating temperature regime than in the constantly warm temperature regime. With EPA supplementation, offspring numbers did not differ significantly between temperature regimes. The increase in offspring production upon EPA supplementation (Fig. 1C, inserted bar graph) and the significant interaction between food and temperature reflect that the positive effect of EPA supplementation on offspring production was higher in the alternating temperature regime.

Population growth rates were also significantly affected by food and temperature (Fig. 1D, Table 1), i.e. they increased upon EPA supplementation in both temperature regimes and were significantly lower in the alternating temperature regime than in the constantly warm temperature regime. The increase in population growth rats upon EPA supplementation (Fig. 1D, inserted bar graph) and the significant interaction between food and temperature indicate that the positive effect of EPA supplementation was higher in the alternating temperature regime than in the constantly warm temperature regime. Mortality was negligible in all treatments.

The body fatty acid composition of all animals was characterized by a number of saturated (mostly C16:0) and monounsaturated (mostly C18:1) fatty acids, which were not significantly affected by temperature and dietary EPA supply (data not shown). Polyunsaturated fatty acids (PUFA) were represented mostly by C18:3n-3 and C18:2n-6. PUFA with more than 18 carbon atoms were not detected in any of the animals, except in those that were reared on the EPA supplemented diet, which contained EPA. C18 PUFA concentrations were significantly affected by temperature but not by food (Fig. 2A, Table 1). Without EPA supplementation, C18 PUFA concentrations were significantly higher in animals exposed to alternating temperatures than in those exposed to the constantly warm temperature. This effect was not significant with EPA supplementation. EPA body concentrations were significantly higher in animals exposed to alternating temperatures than in animals exposed to the constantly warm temperature (ANOVA, F_1,4_ = 20.8, p = 0.01; Fig. 2B). The ratio of saturated to unsaturated fatty acids did not differ significantly among treatments (data not shown).

**Fig. 2:**
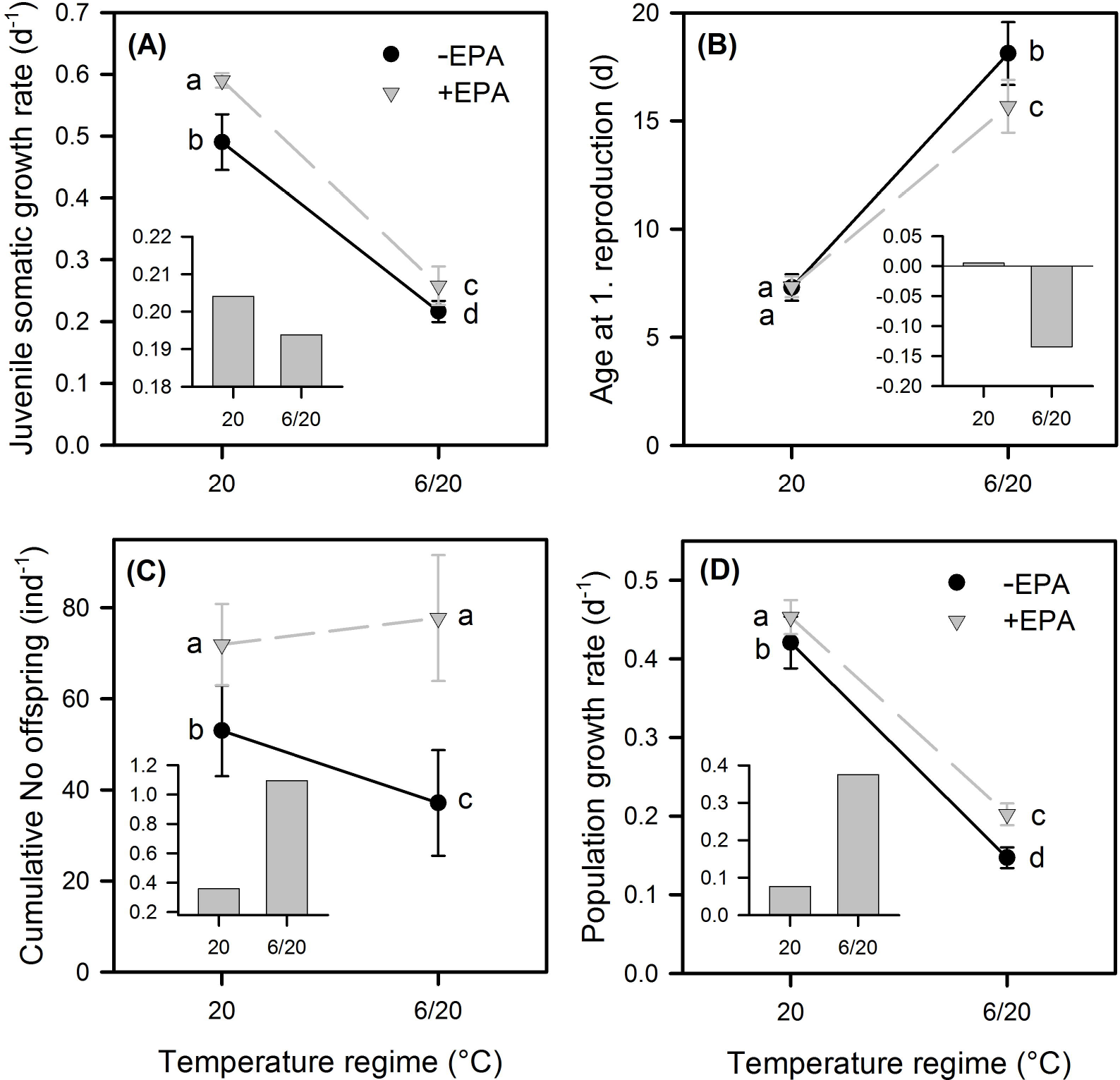
C18 PUFA (A) and EPA (B) concentrations and TBARS equivalents (C) in whole body extracts of *Daphnia magna* exposed to two different temperature regimes, i.e. either constantly 20°C or alternating 6 and 20°C. Data represent means ± standard deviation (n = 3-4). Different letters indicate significant differences among treatments (Tukey’s HSD, P < 0.05). EPA was detected only in animals reared on the EPA supplemented diet.

The concentration of TBARS in whole body extracts was significantly affected by temperature but not by food and did not reveal a significant interaction between temperature and food (Fig. 2C, Table 1). With EPA supplementation, the concentration of TBARS was significantly higher in animals exposed to the alternating temperatures than in those exposed to the constantly warm temperature. This effect was not significant without EPA supplementation.

## Discussion

We show here that the ability of *Daphnia* to cope with alternating temperatures is significantly affected by lipid-mediated food quality. While the effect of EPA supplementation on juvenile somatic growth rates was higher in the constantly warm temperature regime, the effect of EPA supplementation on offspring production and population growth rates was higher in the alternating temperature regime. The more pronounced effect of EPA supplementation on juvenile somatic growth rates in the constantly warm temperature regime presumably reflects that the overall growth potential was higher at constantly warm temperatures and thus also the potential of EPA to increase somatic growth rates. Despite a higher growth potential at constantly warm temperatures, the positive effect of EPA supplementation on offspring production was higher at alternating temperatures. Without dietary EPA supply, offspring production was significantly lower in the alternating temperature regime than in the constantly warm temperature regime. With dietary EPA supplementation, however, offspring production was similar in both temperature regimes, i.e. the negative effect of alternating temperatures on offspring production was equalized by dietary EPA supplementation. This supports previous findings showing that the dietary EPA supply gains in importance at lower temperatures (Sperfeld & Wacker, 2011; Martin-Creuzburg *et al*., 2012). The more pronounced effect of EPA supplementation on reproduction in the alternating temperature regime resulted in higher population growth rates. These findings suggest that vertically migrating *Daphnia*, which in deep temperate lakes are often confronted with similar temperature changes during vertical migration as applied in our study (Stich & Lampert, 1981; Lampert, 2011), rely more on an adequate dietary PUFA supply than non-migrating *Daphnia*. Experiments on the vertical migration behavior of *Daphnia* demonstrated that animals with access to dietary EPA are migrating to deeper water layers, implying that the migration amplitude and thus the capacity to seek shelter from fish predation is affected by the dietary PUFA supply (Brzezinski & von Elert, 2015). Together with our findings, indicating that the ability of *Daphnia* to cope with alternating temperatures increases with dietary EPA supplementation, this suggests that the vertical migration behavior is strongly affected by PUFA-mediated food quality.

*Daphnia* are commonly considered as unselective filter-feeders, which do not discriminate between food particles differing in nutritional quality (DeMott, 1986). The predominance of long-chain PUFA-deficient phytoplankton taxa, such as green algae and cyanobacteria, in the food-rich epilimnion of lakes may constrain the performance of both migrating and non-migrating *Daphnia* populations. However, migrating *Daphnia* populations may be even more susceptible to dietary PUFA deficiencies because the shortage in dietary PUFA may exacerbate the trade-off between light-dependent predation losses and reduced growth and reproduction and thus increase the demographic costs associated with DVM. Our data imply that the vertical migration behavior of *Daphnia* is more pronounced under natural conditions when the seston is dominated by long-chain PUFA-rich algae, such as diatoms or cryptophytes. However, more empirical research is required to assess whether the dietary PUFA supply needs to be considered as another proximate cue triggering DVM (Fig. 3). Moreover, further studies are required to assess the physiological significance of dietary PUFA for DVM in a more natural context, considering that migrating *Daphnia* are typically confronted with rather low food concentrations during the day (Winder, Boersma, & Spaak 2003). Food quality effects often decrease with decreasing food quantity because of changes in nutrient allocation (Sterner & Schulz 1998; Schälicke, Sobisch, Martin-Creuzburg *et al*., 2019). At high food quantity, PUFA can be used to modulate membrane properties or to fuel distinct physiological processes (e.g. the eicosanoid pathway). At low food quantity, however, PUFA might be also catabolized to gain energy. It remains to be tested whether migrating *Daphnia* retain higher amounts of EPA and other PUFA than non-migrating *Daphnia* under natural conditions and whether the positive effect of EPA supplementation observed here is less pronounced when the animals are temporarily exposed to low food quantity.

**Fig. 3:**
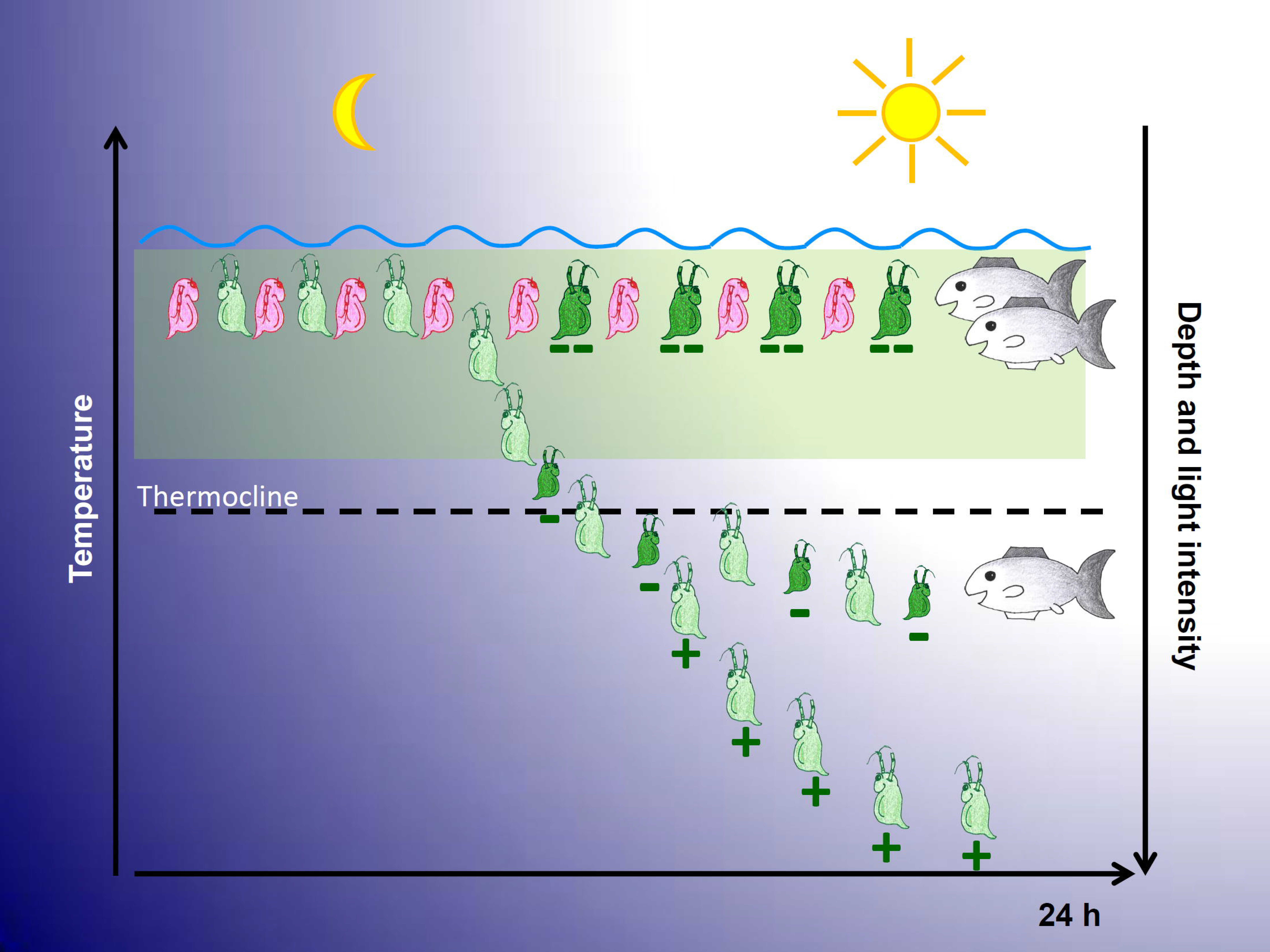
Conceptual model of *Daphnia* diel vertical migration (DVM) considering dietary PUFA supply as an additional proximate factor. In spring, when water temperatures rise and lakes become stratified in a warm and food-rich epilimnion (light green beam) and a cold and food-poor hypolimnion, *Daphnia* start to migrate when fish kairomones are present (Ringelberg & van Gool, 2003). DVM is controlled by the relative change in light intensity, which is greatest in the mornings and in the evenings (Siebeck, 1960). The amplitude of DVM is determined by a variety of different factors, such as water transparency (Dodson, 1990), the concentration of fish kairomones (von Elert & Pohnert, 2000), oxygen availability (Sakwinska & Dawidowicz, 2005), body size (Dini, O’Donnell, Carpenter *et al*., 1987) and potentially the availability of dietary PUFA (Brzeziński & von Elert, 2015). The scheme also shows the potential trade-off between predator evasion and food quality: While non-migrating *Daphnia* remain in the epilimnion during the day (red *Daphnia*), irrespective of dietary PUFA availability, migrating *Daphnia* seek shelter in the hypolimnion given that they are sufficiently supplied with dietary PUFA (green *Daphnia* (+)). The predominance of long-chain PUFA-deficient phytoplankton taxa (green algae, cyanobacteria) may either prevent DVM (green *Daphnia* (--)) or may reduce the migration amplitude (green *Daphnia* (-); see Brzeziński & von Elert, 2015), depending on dietary PUFA concentrations. *Daphnia* that are migrating at moderate dietary PUFA concentrations may be more susceptible to temperature-caused physiological constraints (present study). A shortage in dietary PUFA may thus exacerbate the trade-off between light-dependent predation losses and reduced growth and reproduction and thus increase the demographic costs associated with DVM.

Alternating temperatures presumably impose additional challenges on *Daphnia* in terms of physiological temperature acclimation. Restructuring the lipid composition of membranes is a fundamental mechanism by which ectothermic organisms adapt to changing environmental temperatures. Exposure to low temperatures typically results in an increase in the relative abundance of unsaturated fatty acids in cell membranes, suggesting that these temperature-mediated modifications in membrane fatty acids are crucial to maintain vital membrane properties (Hazel & Williams, 1990; Hazel, 1995). *Daphnia* have limited capacities to modify the incorporation of dietary PUFA and their body fatty acid profile largely reflects that of their food (Brett, Müller-Navarra, Ballantyne *et al*., 2006). There is evidence, however, that *Daphnia* accumulate higher amounts of PUFA within their body when exposed to colder temperatures (Schlechtriem, Arts, & Zellmer, 2006); whether this also involves a higher degree of fatty acid unsaturation within membrane phospholipids remains unclear. In our experiment, *Daphnia* exposed to alternating temperatures contained higher amounts of C18 PUFA than animals exposed to constantly warm temperatures. However, this effect was significant only without EPA supplementation. In animals exposed to the alternating temperatures, EPA supplementation resulted in significantly lower body C18 PUFA concentrations. Instead, animals exposed to the alternating temperatures contained significantly higher amounts of EPA, suggesting that the supplemented EPA was used as a substitute for C18 PUFA. The concentration of EPA within the body was generally much lower than that of the predominant C18 PUFA, 18:3n-3 and 18:2n-6, which reflects dietary availability. Assuming that EPA was used as a substitute for C18 PUFA implies that EPA is more efficient in modulating membrane-related temperature acclimation than C18 PUFA, which has been suggested previously based on experiments showing that the positive effects of EPA supplementation on a C18 PUFA-rich diet is higher at colder temperatures (Martin-Creuzburg *et al*., 2012).

Higher body PUFA concentrations in animals exposed to the alternating temperatures are in accordance with the homeoviscous adaptation hypothesis, considering that the animals in the alternating temperature regime were on average exposed to colder temperatures. However, exposure to alternating temperatures may also hamper lipid-mediated temperature acclimation. Despite the significance of PUFA for growth and reproduction, high body PUFA concentrations may also become detrimental because PUFA are sensitive to lipid peroxidation, resulting in radical formation and oxidative stress (Esterbauer, Schaur, & Zollner, 1991; Yin *et al*., 2011). The influence of temperature on oxidative stress is controversial, albeit most studies on ectothermic animals revealed increasing oxidative stress with increasing temperature (Lushchak, 2011). In *Daphnia*, lipid peroxidation has been found to increase in response to heat exposure, but did not appear to be a predictive measure of susceptibility to thermal damage (Coggins, Collins, Holbrook *et al*., 2017). To the best of our knowledge, lipid peroxidation in response to fluctuating temperatures has not been studied yet. We show here that the concentration of thiobarbituric acid (TBA)-reactive substances (TBARS), as a measure of the major lipid peroxidation end product malonic dialdehyde (MDA), is higher in animals exposed to fluctuating temperatures than in animals exposed to constantly warm temperatures, suggesting that exposure to fluctuating temperatures resulted in increased lipid peroxidation. Although this effect was observed with and without EPA supplementation, it was significant only for the EPA supplemented diets, suggesting enhanced temperature-mediated lipid peroxidation in the presence of long-chain PUFA. It has been proposed that lipid peroxidation in *Daphnia* may increase in response to heat exposure as a by-product of changes in phospholipid fatty acid unsaturation (Coggins *et al*. 2017). In our study, higher EPA concentrations in *Daphnia* exposed to fluctuating temperatures were accompanied by higher lipid peroxidation, implying a trade-off between long-chain PUFA accumulation and oxidative stress during DVM. The accumulation of long-chain PUFA as a mechanism to cope with low temperatures experienced during the day may thus come at the expense of increased oxidative stress associated with high temperature exposure during the night. Our data suggest, however, that the positive effects of EPA supplementation outweigh potential negative effects associated with lipid peroxidation during DVM.

## Conclusions

We show here that the well-known positive effect of dietary EPA supplementation on offspring production and thus population growth rates of *Daphnia* is more pronounced at alternating temperatures than at constantly warm temperatures, suggesting that an adequate dietary PUFA supply is important especially for migrating *Daphnia*. Exposure to alternating temperatures caused modification in body PUFA concentrations and, consequently, increased lipid peroxidation. Detrimental effects of lipid peroxidation on *Daphnia* performance were not evident, suggesting that the positive effects of PUFA accumulation at alternating temperatures mask potential negative effects of PUFA-mediated oxidative stress. Our data imply that a dietary deficiency in long-chain PUFA has strong implications for predator evasion and thus *Daphnia* population dynamics.

## Acknowledgements

We thank M. Wolf and P. Merkel for excellent technical assistance and A. Wacker for statistical advice and support.

## Author Contributions

DMC designed the experiment. MF conducted the experiment and performed the lipid analyses. BZ performed the lipid peroxidation assay. All authors analysed the data. JIN and DMC wrote the first draft of the manuscript. All authors contributed to subsequent versions of the manuscript.

## Data accessibility

The datasets used and/or analysed during the current study are available from the corresponding author on reasonable request.

